# Eye movements during visuomotor adaptation represent only part of the explicit learning

**DOI:** 10.1101/724864

**Authors:** Zohar Bromberg, Opher Donchin, Shlomi Haar

## Abstract

Visuomotor rotations are learned through a combination of explicit strategy and implicit recalibration. However, measuring the relative contribution of each remains a challenge and the possibility of multiple explicit and implicit components complicates the issue. Recent interest has focused on the possibility that eye movements reflect explicit strategy. Here we compared eye movements during adaptation to two accepted measures of explicit learning - verbal report and the exclusion test. We found that while reporting, all subjects showed a match between all three measures. However, when subjects did not report their intention, the eye movements of some subjects suggested less explicit adaptation than what was measured in an exclusion test. Interestingly, subjects whose eye movements did match their exclusion could be clustered into two subgroups: fully implicit learners showing no evidence of explicit adaptation and explicit learners with little implicit adaptation. Subjects showing a mix of both explicit and implicit adaptation were also those where eye movements showed less explicit adaptation than did exclusion. Thus, our results support the idea of multiple components of explicit learning as only part of the explicit learning is reflected in the eye movements. Individual subjects may use explicit components that are reflected in the eyes or those that are not or some mixture of the two. Analysis of reaction times suggests that the explicit components reflected in the eye-movements involve longer reaction times. This component, according to recent literature, may be related to mental rotation.

**Significance Statement:** Visuomotor adaptation involves both explicit and implicit components: aware re-aiming and unaware error correction. Recent studies suggest that eye movements could be used to capture the explicit component, a method that would have significant advantages over other approaches. We show that eye movements capture only one component of explicit adaptation. This component scales with reaction time while the component unrelated to eye movements does not. Our finding has obvious practical implications for the use of eye movements as a proxy for explicit learning. However, our results also corroborate recent findings suggesting the existence of multiple explicit components, and, specifically, their decomposition into components correlated with reaction time and components that are not.

## Introduction

Visuomotor adaptation is commonly used to study human motor learning in health (e.g. Galea et al., 2015; Ghilardi et al., 2000; Haar et al., 2015a; Krakauer et al., 1999; Taylor and Ivry, 2013) and disease (e.g. Rabe et al., 2009; Wong et al., 2019). In a visuomotor rotation task, the visual representation of hand position is manipulated such that subjects must learn a new mapping of motor commands to apparent outcomes. Recent studies dissociated explicit and implicit processes in the visuomotor adaptation task (Hegele and Heuer, 2010; Mazzoni and Krakauer, 2006; Taylor and Ivry, 2011; Taylor et al., 2014; Werner et al., 2015), where the sum of the two gives the total adaptation.

One measure of implicit learning is to ask subjects to reach straight for the target without perturbation (or without any visual feedback) and to measure the difference between the direction of reach and the target. We call this exclusion because the subject is being asked to “exclude” their explicit knowledge from their behavior. When measured after adaptation, this has also been called the after-effect. When measured during adaptation, it is sometimes called a ‘catch trial’ (Werner et al., 2015). By its nature, exclusion cannot be measured every trial since it presumes surrounding adaptation trials. In order to assess implicit and explicit learning throughout the adaptation process, Taylor et al., (2014) suggested simply asking subjects to report aiming direction before each movement by reporting which of the numbers displayed in a circle on the screen was in the direction the subject intended to move. Reporting has been a very productive experimental approach. However, the protocol has known limitations, e.g., reporting increases the length and variability of inter-trial intervals since subjects can start moving only after reporting.

One alternative is to measure explicit learning using eye-movements: perhaps eye movements can provide an objective measure of subjects’ intentions without needing special trials or direct questioning. During unperturbed reaching movements, the eyes were found to provide an unbiased estimate of hand position (Ariff et al., 2002). During visuomotor rotation there is an increase correlation between gaze and hand directions in early practice which gradually decreased thereafter (Rand and Rentsch, 2016). Indeed, a recent study found that individual differences in gaze patterns during visuomotor adaptation were linked to participants’ explicit learning (de Brouwer et al., 2018). Interestingly, they noticed subjects whose eye movements did not reflect adaptation while their aftereffects did indicate some explicit learning. Thus, raises the possibility that some forms of explicit adaptation are captured by the eye movements, while others are not.

This possibility is in line with recent suggestions of multiple explicit strategies in human motor learning, even in a redundant visuomotor rotation task (McDougle and Taylor, 2019). McDougle and Taylor interpret their results to show that subjects in different conditions may use either discrete response caching or parametric mental rotation as two different explicit strategies. Their results further suggest that reaction time (RT) can be used to dissociate these explicit strategies: mental rotation is a time-consuming computation and caching is a fast automatic process that does not require a long RT (Haith and Krakauer, 2018). Here we set to explore the different explicit components captured or not by eye movements, and their link to the explicit strategies captured by RT.

In the first experiment of the current study, we measured subjects’ eye movements during visuomotor rotation with verbal reporting and without. Like in de Brouwer et al. (2018), our results demonstrate that, in verbal reporting, eye fixations before movement onset accurately predicts the reported aiming direction. Without reporting, eye fixation before movement onset correlates well with explicit learning measured by after effect. However, it does not account for the full explicit knowledge revealed by exclusion. This suggests that only a component of explicit learning is being captured by eye movements when there is no verbal report.

In a second experiment we explore the time course of the discrepancy between eye movements and exclusion by introducing exclusion (catch) trials in addition to testing for an after effect. For some subjects, measures of explicit learning from eye movements matched those from exclusion. For other subjects, exclusion revealed more explicit knowledge than that found in the eye movements. The first group divided into two subgroups: those using primarily explicit strategy and those with hardly any contribution from an explicit strategy. The second group, where exclusion showed more explicit knowledge than did the eye movements, showed subjects with the full range of combinations of explicit and implicit learning. Further analysis of RT seems to indicate that the explicit knowledge reflected in the eye movements may be the same mental rotation component identified by McDougle and Taylor (2019).

## Methods

### Participants

114 right-handed subjects with normal or corrected-to-normal vision participated in the study: 44 subjects (31 female, aged 18-29) in the first experiment, and 70 subjects (46 female, aged 18-31) in the second experiment. All subjects signed an informed consent form which also asked for basic information about their relevant medical status. None of the subjects reported neurological or motor impairments. The experimental protocol was approved by the HSR (Human Subject Research) committee of Ben-Gurion University of the Negev.

### Apparatus

Participants were comfortably seated on a height-adjustable chair at a table with a digitizing Wacom Intuos Pro tablet with an active area of 311 × 216mm and a polling rate of 2154 Hz. They faced a 19” vertical computer screen. Their eye-movements were recorded with an EyeLink 1000 (SR Research) eye-tracking system, with sampling rate of 500Hz and the accuracy of approximately 0.5°. The participants rested their head on a stabilizing support which included braces for the chin and forehead. Calibration of the eye tracking system was done for each subject before the experiment. During the experiment, subjects made center-out, horizontal reaching movements on the surface of the tablet using a digital stylus. On the monitor, they saw a cursor and targets to move to (detailed in the next section). The cursor’s movements matched the movements of the tablet pen (except as detailed in the next section). The eye link system provides the location of the pupil, and timing and location of events including blinks, fixations and saccades. In the reporting group of the first experiment, the number subjects reported as their intended aiming direction were recorded and typed into an excel sheet (details below).

### Trial structure

We used three different trial types in our experiments. The general structure of these trial types is shown in Figure 1A. At the beginning of every trial, subjects moved a grey cursor to a white origin. The origin appeared in the center of the screen and corresponded to a hand position at the center of the tablet. After maintaining this position for 1000 ms, a green target circle appeared at a distance of 8 cm on the screen (corresponding to a movement of 5.5 cm on the tablet) and visual landmarks appeared surrounding the target. Targets could be in one of the 8 cardinal and intercardinal directions relative to the origin. The order of targets was pseudo-random for each subject and between subjects such that a complete circuit of the targets was completed every 8 movements. The cursor disappeared when the participant’s hand was further than 0.56 cm from the center of the origin. When the hand crossed the distance of 7.6 cm (95% of the target distance) from the origin, a red circle the same size as the original cursor appeared on the screen at the same distance as the target. This circle was presented in order to give subjects visual feedback about the cursor position at the end of the trial. The red circle that indicated the cursor location remained visible for 350, 700, or 1000 ms. Different presentation times were used in different groups and in different experiments. After the red circle disappeared a white ring appeared centered at the origin with radius equal to the distance of the hand from the origin. This ring guided the hand back to the origin without providing information about its exact location.

**Figure 1.**
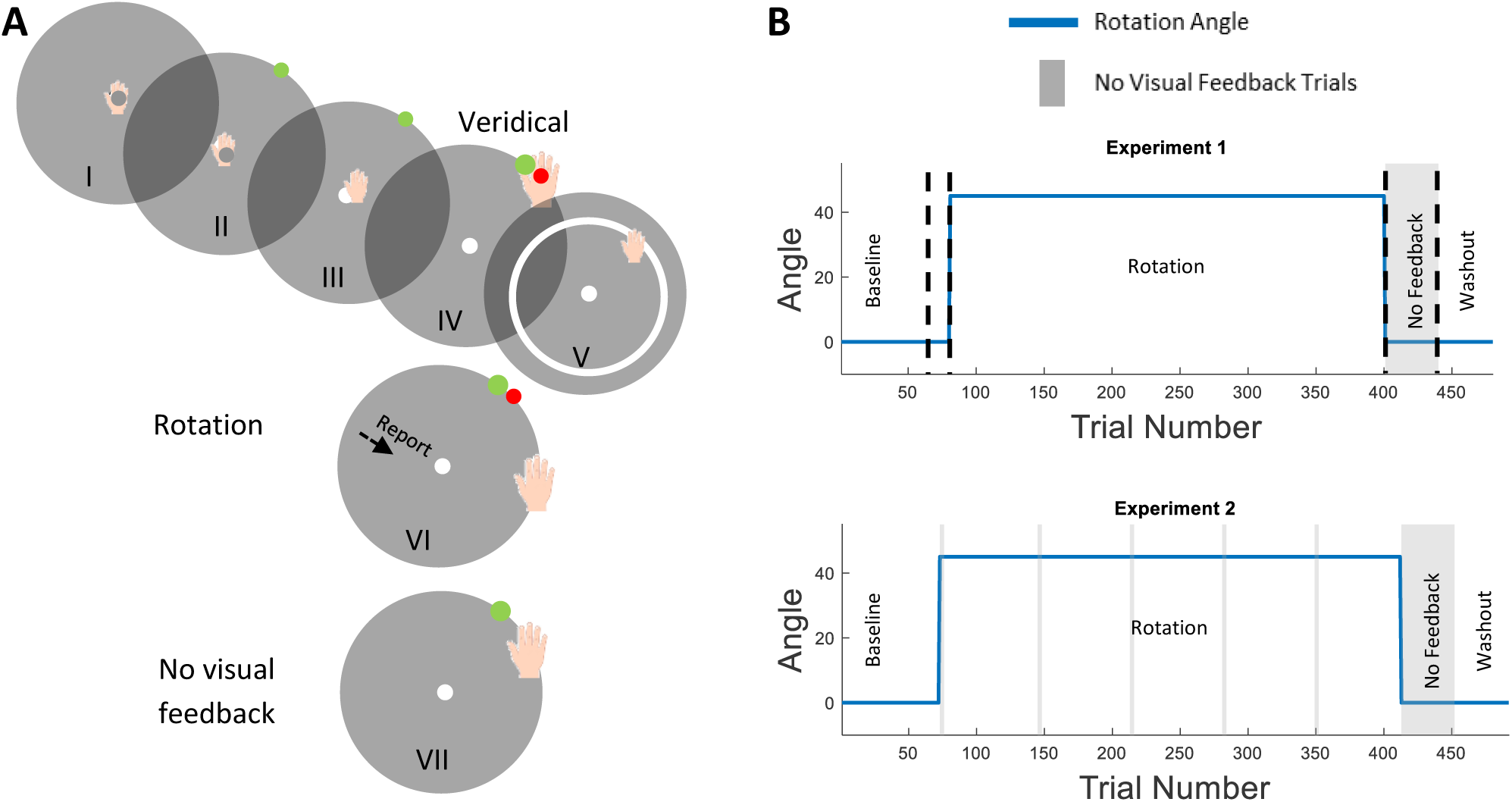
Trial and Experiment design. (**A**) Basic trial structure. (I) Subjects performed reaching movements from the origin. (II) After one second, a target appeared, and (III) subject started reaching towards the target. (IV) On veridical trials, when reaching to target, a red circle appeared indicating the hand position. (V) After the feedback disappeared, a white ring appeared, which directed the hand back to the origin. (VI) In rotation trials, the red circle appeared rotated 45° counterclockwise. In the report condition, subjects were asked to report before initiating their hand movement. (VII) In no visual feedback trials, subjects received no feedback at the end of the movement. (**B**) Experimental design. Top: First experiment. In the first and second Baseline Block, feedback was veridical. In the Rotation Block, the cursor was rotated by 45°. In the No Visual Feedback Block, the landmarks and cursor feedback were removed, and participants were instructed to aim directly to the target. In the Washout Block, conditions were similar to the first Baseline Block. In the Second Baseline Block and in the Rotation Block, participants in the R condition reported their aiming direction. Bottom: Second Experiment. Similar to the first experiment, however, without the second Baseline Block and without report sessions. In addition, five Mini Exclusion Blocks were spread throughout the Rotation Block.

The three different trial types were: *Veridical feedback*, the red circle appearing at the end of each trial reflected the true location of the hand. *Rotation*, the red circle appeared at a location that was rotated by 45° counterclockwise relative to the hand movement. *No visual feedback*, no red circle appeared at all, nor did any other landmarks. In no visual feedback trials, subjects were instructed to aim straight towards the target. In all trials, subjects received auditory feedback to control movement speed: movements that reached a distance of 7.6 cm from the origin center within 500 ms were rewarded with a pleasant “ding” sound; otherwise, subjects heard an unpleasant “buzz” sound.

### Data collection

x and y coordinates of the hand trajectories were collected by the tablet and saved from the moment of target appearance until the hand reached 95% of the targets circle radius. Eye movements were recorded continuously using the eye tracker. Eye movement data was pre-processed to remove blinks (as recorded by the eye tracker software). In addition, eye movements were re-centered, to correct for drift over the experiment, by assuming that the eye is fixated on the origin during the 1000 ms before the hand movement begins. Re-centering was accomplished by accumulating eye position in this time window during the current trial, two preceding trial and two following trials, and taking the median position across the entire 5000 ms of data.

### Experiment 1

#### Groups

This experiment had two groups of 22 subjects each: Report and No-Report. Each subject was assigned to one of two feedback times: 350ms or 700 ms, counterbalanced between the groups. The feedback time showed no effect on task performance, learning, and eye movements in either of the groups, and thus we did not address this subdivision in the results.

#### Procedure

Each trial presented one of the eight targets in cardinal and intercardinal directions (0°, 45°, 90°, 135°, 180°, 225°, 270°, 315°) relative to the origin. For subjects in the Report group, 63 numbered landmarks spaced by 5.625° appeared on a circle with radius 8 cm around the origin (this is the same radius at which targets appeared). Low numbers were nearer the target. Before each movement, subjects were instructed to say out loud the number towards which they were aiming towards in order to get the cursor to the target. For subjects in the No-Report group, hollow circles were presented instead of the numbered landmarks and they were not asked to report their intended aiming direction. Indeed, the No-Report group was not informed in any way that they might want to aim to a direction different than the target. The experimental sequence was the same for both groups. Each session was divided into 5 blocks: two baseline blocks (72 and 8 trials) consisting of veridical trials. The first baseline (*veridical feedback*) block allowed subjects to get familiar with the reaching task and the second block was intended for subjects in the Report group to practice the report. For subjects in the No-Report group, there was no difference between these two blocks. In the third block, the rotation block (320 *rotation* trials), the cursor was rotated relative to the origin. Subjects in the Report group were required to report their aiming direction during this block. The fourth block was a no feedback block (40 trials), which consisted of *no visual feedback* trials. In the last block, a washout block (40 trials), subjects were presented with *veridical feedback* trials (Figure 1B). The percentage of successful trials was displayed at the end of each block. A trial was considered successful if the red circle was within 5° of the target. Every 40 trials, the experimental program displayed a full-screen text message reminding subjects of the instructions.

### Experiment 2

#### Procedure

Each trial presented one of the eight targets in secondary intercardinal directions (22.5°, 67.5°, 112.5°, 157.5°, 202.5°, 247.5°, 292.5°, 337.5°) relative to the origin. In this experiment, trial feedback was visible for 1,000 ms. 47 hollow circles spaced 7.5° apart appeared on the target circle as landmarks surrounding the target before each movement. The experiment was divided to 4 blocks: a baseline block with 72 *veridical feedback* trials, a rotation block with 320 *rotation* trials, a no feedback block with 40 *no visual feedback* trials, and the last block, a washout block with 40 *veridical feedback* trials. 20 more *no visual feedback* trials were presented during the rotation block (Figure 1B). They were evenly spaced during the block in 5 mini blocks of 4 trials each. Subjects were instructed at the beginning of the experiment that their goal is to hit the target with the cursor. All instructions that were given during the experiment were presented on the screen. After the first 2 trials of the rotation block, a message appeared on the screen asking the subject to pay attention to the error and to hit the target with the cursor. In addition, before and after each mini block of *no visual feedback* trials, a message appeared announcing the beginning and end of this block. In the beginning, the message instructed subjects to ignore their strategy and to hit the target. In the end, the message instructed them to go back to using their strategy.

### General Data analysis and statistics

#### Hand movement analysis

The Hand-Target Difference was calculated as the difference between the target location and the hand position when the cursor reached 95% of the distance from origin to the targets. Trials, in which the movement from the origin towards the target was not strictly increasing after the cursor passed 5% of the distance to the targets or trials in which movement was too slow, were excluded. In the first experiment, subjects were presented with text reminding them of the instructions every 40 trials. Trials immediately following these reminders were discarded, since subjects often tested the degree of rotation by aiming directly to the target. Each movement RT was defined as the time between target appearance and the cursor reaching 7% of the target distance.

#### Eye movement analysis

The eye movements before movement initiation followed a stereotypical pattern: during the baseline block, subjects fixated first on the origin and then on the target. During the rotation block, target fixation was often followed by eye movements that carried the gaze in the direction opposite to the rotation (Figure 2). We tested several measures of this latter gaze shift to see which best correlated with subjects’ reported aiming direction. All methods produced similar results and the choice of measure did no influence any of our findings. Thus, following previous results showing the eye leading upcoming hand movements with a similar constant lead (Ariff et al., 2002), we choose to use the last fixation before movement onset to characterize subjects’ intended aiming direction and called it the Explicit Eye. If eye fixation before movement onset was missing (due to a blink), or if it was near the origin (radius<50% of the target radius), or beyond the target area (radius>150% of the target radius) the eye movements for that trial were discarded. Any subject for whom over 50% of the rotation trials were discarded were excluded from further analysis (4 in the No-Report group in the first experiment, and 1 in the second experiment). We used the term Implicit Eye for the difference between the Explicit Eye and the Hand-Target Difference; it represents the estimated implicit adaptation derived using the eye movements.

**Figure 2.**
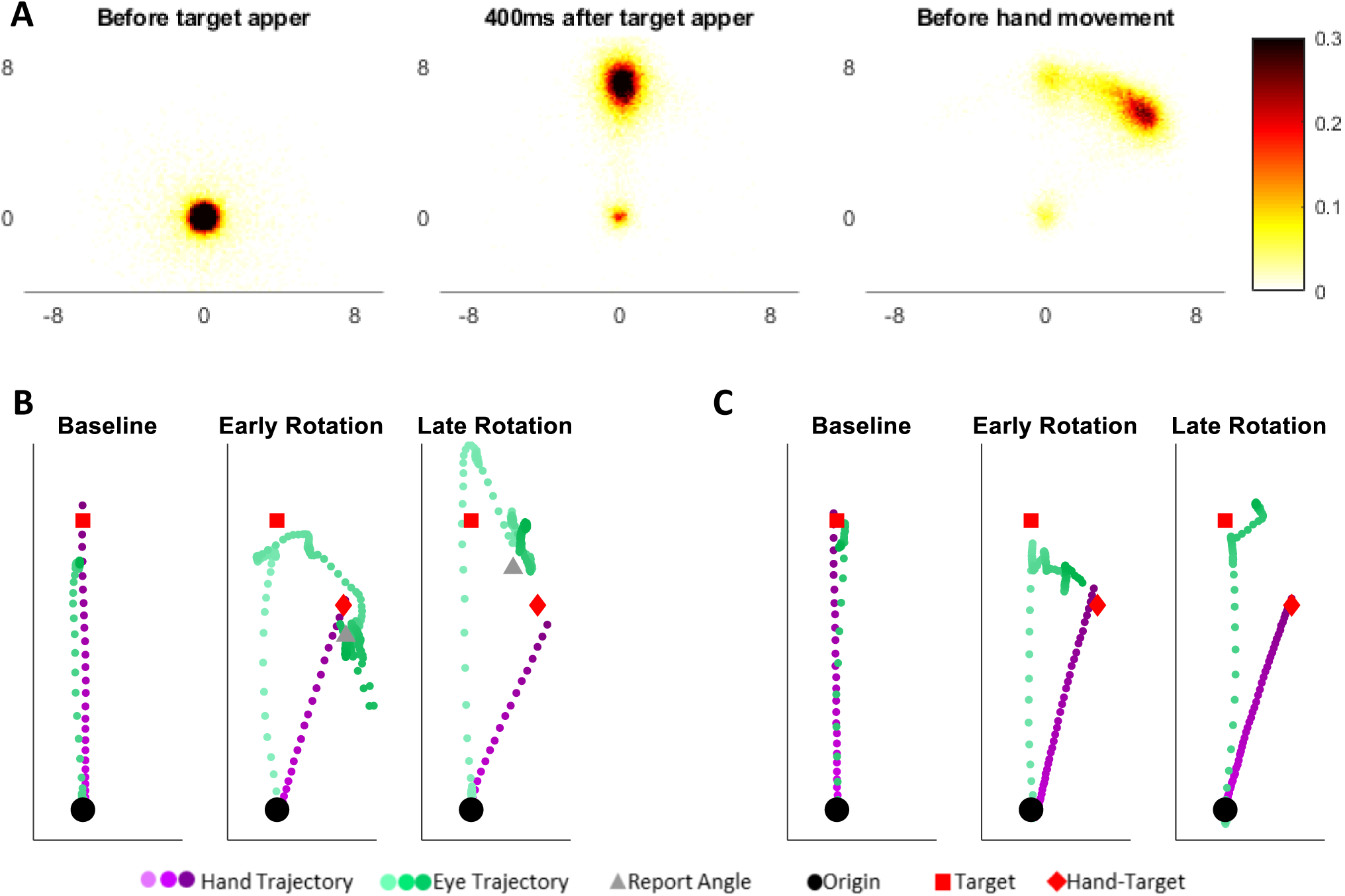
Eye movements pattern. (**A**) Fixations distributions during the phases of the trial prior to movement initiation, in early rotation trials. (**B+C**) Hand and eye trajectories of two individuals from the Report (B) and the No-Report (**C**) group. In the Baseline Block, the subjects first gazed towards the target and later made a reaching movement towards it. At the beginning of the Rotation Block, both subjects shifted their gaze toward the target and later toward the hand target, the reporting subject then reported a direction close to the hand-target, and both subjects moved their hand toward the hand-target. By the end of the rotation, the subjects also shifted their gaze first to the target, then toward the side opposite of the rotation. However, this secondary shift was smaller than the hand target angle for the reporter and even smaller for the non-reporter. The reporter also reported a smaller angle. Both kept moving their hand moved to the hand-target direction.

#### Reporting analysis

Report trials (in the first experiment) were also characterized by the subject’s statement of their intended aim direction. The reported number was multiplied by 5.625° / landmark in order to convert it to an aiming angle. This was called the Explicit Report. The Implicit Report was calculated by subtracting the Explicit Report from the Hand-Target Difference. This is the difference between where subjects said they aim and where they moved their hand.

#### No visual feedback trials analysis

In the *No Visual Feedback* trials, subjects were instructed to aim directly towards the target. Thus, the difference between the hand and the target in these trials represents, by definition, residual implicit knowledge of the rotation not under the subject’s control. In the second experiment, there were 20 *no visual feedback* trials in mini blocks during the rotation block, and a no visual feedback block after the rotation block. The mini blocks trials and the first four trials of the no visual feedback block were named the Implicit No Visual Feedback. We excluded 3 subjects who later reported they didn’t understand the instruction during those trials.

#### Statistics

Unless otherwise noted, we report the mean and 95% confidence intervals (in brackets) for all reported and depicted values. A non-parametric bootstrap with 10000 samples was used to generate sampling distribution for each group means. We took the percentiles of the sampling distribution to get the confidence interval. We forego reporting statistical significance as per the recommendations from Amrhein et al. (2019). Calculations of traditional hypothesis tests and effect sizes can be found in the supplementary material. In addition, all data and code for the data analysis are available online at: osf.io/dsn3v.

The measures defined for each subject and each trial were the difference between the hand and the target (Hand-Target Difference), the last fixation before movement onset (Explicit Eye) and the RT. In the first experiment, the report group also has the reported angle of the aim direction (Explicit Report). For each measure, we smoothed the results by averaging over bins of eight sequential trials (one for each target). For each bin, trials that were more than two standard deviations from the mean of the bin were removed (in the first experiment 2.0% of the available Hand-Target Difference trials were outliers and 2.3% from the available Explicit Eye trials, in the second experiments 1.9% and 2.3% trials were removed, respectively). We then calculated the implicit measures for each trial by subtracting the corresponding explicit measures (Eye and Report) from the Hand-Target Difference to create the Implicit Eye and Implicit Report. In case of missing values in a trial in Hand-Target Difference, Explicit Eye or Explicit Report (due to outliers, slow movements, blinking, missed report, etc.), no implicit measure was calculated for that measure for that trial. For each measure, we used a parametric bootstrap to find the sampling distribution of the mean for each bin, resampling data from a normal distribution with the outlier corrected mean and standard deviation determined by the eight points in each bin for each measure (or fewer points if outliers were discarded).

In the second experiment, we also measured Implicit Exclusion. We averaged over the four sequential trials of each mini-block to create a binned version of Implicit Exclusion, and again used a parametric bootstrap to find the sampling distribution of the mean of each bin, resampling data from a normal distribution with mean and standard deviation determined by the four points in each bin. Similarly, we also averaged over the Hand-Target Difference, the Explicit Eye, and the Implicit Eye in the four trials preceding each Mini Exclusion Block. The Explicit Exclusion was calculated by subtracting the binned Implicit Exclusion from the binned Hand-Target Difference before the Mini Exclusion Block. Implicit difference was defined by subtracting the binned Implicit Eye before the Mini Exclusion Block from Implicit Exclusion. Since for some subjects some bins were missing, we used PPCA (Tipping and Bishop, 1999) to fill in the missing values by projecting the data (6 bins per subject) down to the five principal components (PCs) space, and then with the PPCA coefficients projecting back to the original 6 dimensional space.

In the reaction time analysis, we binned the Hand-Target Difference and the Explicit Eye according to the RT. We divided the reaction time into windows of 25 ms from 0 ms to 2000 ms and averaged over the Hand-Target Difference and the Explicit Eye of all trials with RT in the same window. The SEM for each window was calculated by dividing the standard deviation of each bin by the square root of the number of trials in this bin.

In all learning curves we focused on three phases: the ‘initial rise’ was defined as the 3rd bin in the rotation block; the ‘late early rise’ was defined the 10th bin in the rotation block (after 80 *rotation* trials), and the ‘end of adaptation’ was defined as the last bin of the rotation block.

#### Clustering

In the second experiment, we clustered the subjects with fuzzy c-means (FCM) clustering. We tested clustering with between two and six clusters. Following Haar et al. (2015b) we used a cluster validity index proposed by Zhang et al. (2008). This index uses a ratio between a variation measure in each cluster and a separation measure between the fuzzy clusters. The smaller the ratio, the better the clustering. Clustering was applied in two steps: The first was on the difference between the two measures of implicit in each of the 6 mini-blocks: the difference between Implicit Eye before the *no visual feedback* trials and the hand at the *no visual feedback* trials. The second step was applied on the Explicit Eye during rotation trials: For each cluster found in the first step, perform additional clustering on the first 3 principal components from the 40 bins of rotation trials.

## Results

We recorded eye movements that subjects made during hand reaching movements perturbed by a clockwise visuomotor rotation to study the relation between subjects’ gaze and explicit learning. In the first experiment, we developed and validated our Explicit Eye measure, and, in the second experiment, we explored the time course of the Explicit Eye measure and its relation to exclusion, a measure of implicit learning.

### Experiment 1

The first experiment followed the protocol of Taylor et al., (2014) and compared reporting (R) and non-reporting (NR) learning groups. The averaged learning curves for both groups showed a rapid rise in the Hand-Target Difference (Figure 3A&B). The initial rise (the 3rd bin in the rotation block) was 32.9 ± 6.2° and 24.7 ± 8.5° for the R and NR groups respectively. The differences between the groups in the initial rise was 8.2 ± 10.5°. Adaptation does not saturate after the initial rise but continues more slowly. This second time constant continues until subjects have zero error. The report group reached adaptation plateau after about 80 trials with mean Hand-Target Difference of 46.5 ± 2.8° at the late early rise. The NR group was slower to reach plateau and at the late early rise their mean Hand-Target Difference was only 33.9 ± 5.4°. The difference between groups in this late early adaptation phase is 12.6 ± 6.1°. By the end of adaptation phase, the mean Hand-Target Difference is 45.8 ± 2.1° and 43.0 ± 2.7° in the R and NR groups respectively.

**Figure 3.**
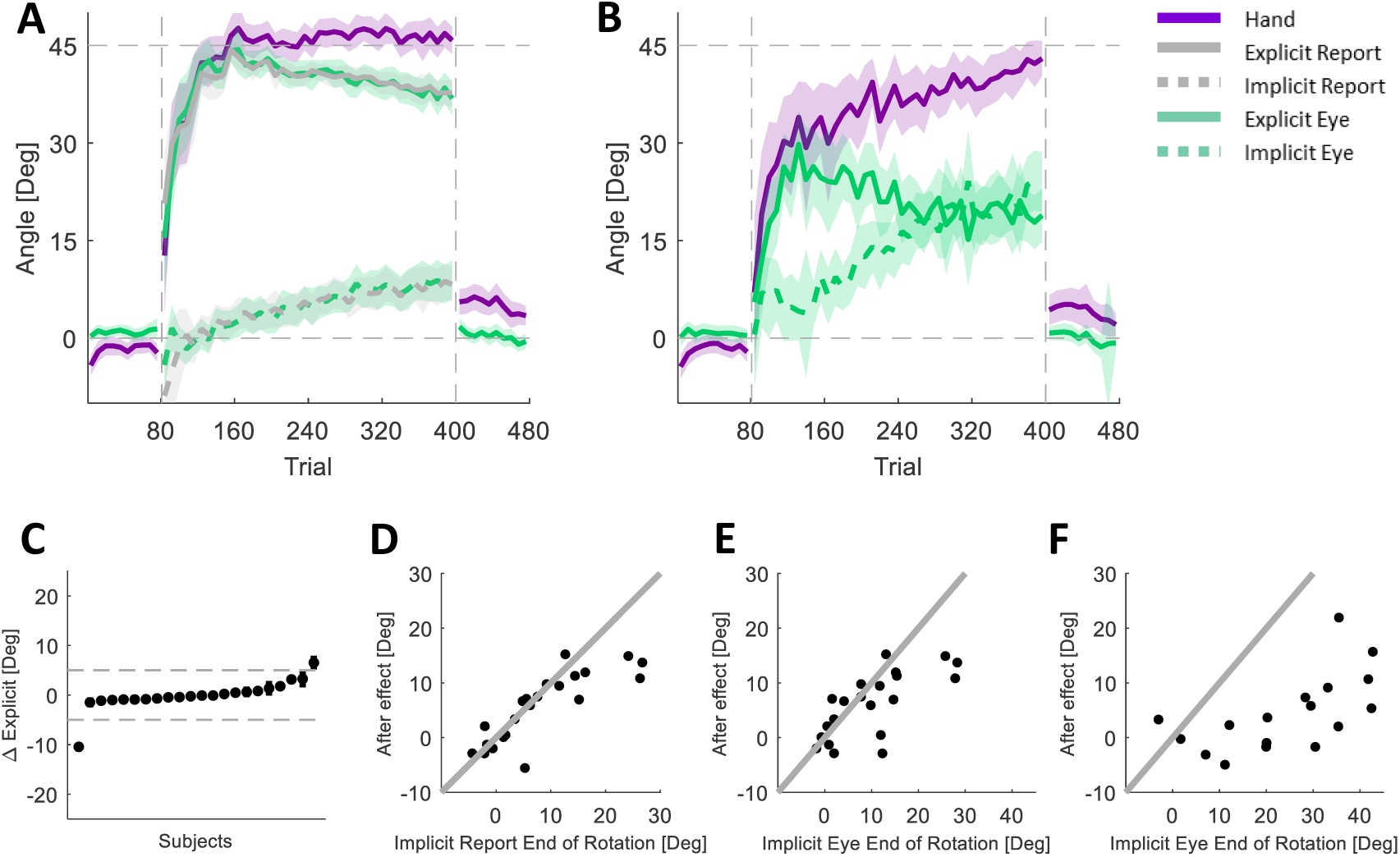
Experiment 1. (**A**) Learning curves of the Report group. (**B**) Learning curves of the No-Report group. (**C**) Subjects’ difference between the Explicit Report and the Explicit Eye. Each dot is the averaged difference between the Explicit Report and the Explicit Eye during the Rotation Block, and each line shows the SEM of this difference. Dashed gray lines show 5° rope. (**D**) The relation between the after effect and the Implicit Report at the end of adaptation in the Report group. (**E**) The relation between the after effect and the Implicit Eye at the end of adaptation in the Report group. (**F**) The relation between the after effect and the Implicit Eye at the end of adaptation in the No-Report group. (D-F) Dots denote individual subjects, identity line in black.

Explicit Report and Explicit Eye in both groups rise quickly and fall off slowly, reflecting a slow but steady increase in implicit knowledge. Differences between groups in the Explicit Eye at the initial rise (difference of 15.8 ± 8.6°) and in the late early rise (19.5 ± 6.0°) mirrors the difference in the Hand-Target Difference, reflecting the dominant role of explicit knowledge in initial learning across both groups. By the end of adaptation, the differences between R and NR still exist (18.0 ± 4.8°).

In the R group, Explicit Eye and Explicit Report match with an average difference of - 0.02 ± 0.8°. This is true not only on average but also for each subject (Figure 3C). As a consequence, the corresponding implicit measures also match. The average difference is −0.3 ± 0.8°, and this is consistent with the aftereffect measure of the implicit: the difference between the after effects and the implicit components are 3.4 ± 3.0° (Figure 3D&E). This contrasts with the consistent lack of a match between Implicit Eye and aftereffect in the NR group (Figure 3F). For the NR group, the Implicit Eye measure suggested a much larger implicit component than revealed by the aftereffect, where the difference between the aftereffects and the Implicit Eye were 19.6 ± 4.5°. Nevertheless, the aftereffect and the Implicit Eye correlated in both groups, with Spearman correlations of 0.75 ± 0.09 and 0.67 ± 0.11 for the R and NR groups respectively. Hence, though the Implicit Eye is higher than the aftereffect in the NR group, they are still correlated.

### Experiment 2

While in the report group the eye movements fully reflected the explicit component of adaptation, in the NR there was a gap. To explore this gap, we conducted a second experiment where we added exclusion trials during the rotation block. Those trials were *no visual feedback* trials in which we asked subjects to ignore their strategy and aim directly at the target. Since, by aiming at the target, they remove the expression of their explicit knowledge, the remaining Hand-Target Difference reflects only their implicit learning. Five mini-blocks of 4 exclusion trials were used during the rotation and these were combined with a virtual mini block of the first four exclusion trials of the aftereffect. This led to a total of 24 exclusion trials in six mini blocks.

Like in the NR group of the first experiment, the Hand-Target Difference initially rose quickly, and later continued to rise slowly (Figure 4A). In this experiment, on average, subjects did not reach full adaptation. Here, too, the Explicit Eye rises initially quickly. However, it did not decline as much as in the first experiment. The Implicit Eye also rose slowly, but less than in the first experiment. (This is a direct result of the decline of the Explicit Eye mentioned above because the Implicit Eye is simply the difference between Hand-Target Difference and Explicit Eye). Finally, here too, there was a gap between the Implicit Exclusion and the Implicit Eye. On average, the Hand-Target Difference showed initial rise (by the 3^rd^ bin) of 25.2 ± 3.7°. By the end of adaptation, the Hand-Target Difference reached 40.2 ± 2.2°. The Explicit Eye, on average, had initial rise of 20.2 ± 3.1° and stayed at about the same level until the end of the adaptation, where it reached 23.3 ± 2.7°. The Implicit Exclusion was lower than the Implicit Eye with a bias of 5.4 ± 1.3°that was not, however, consistent across subjects (Figure 4B).

**Figure 4.**
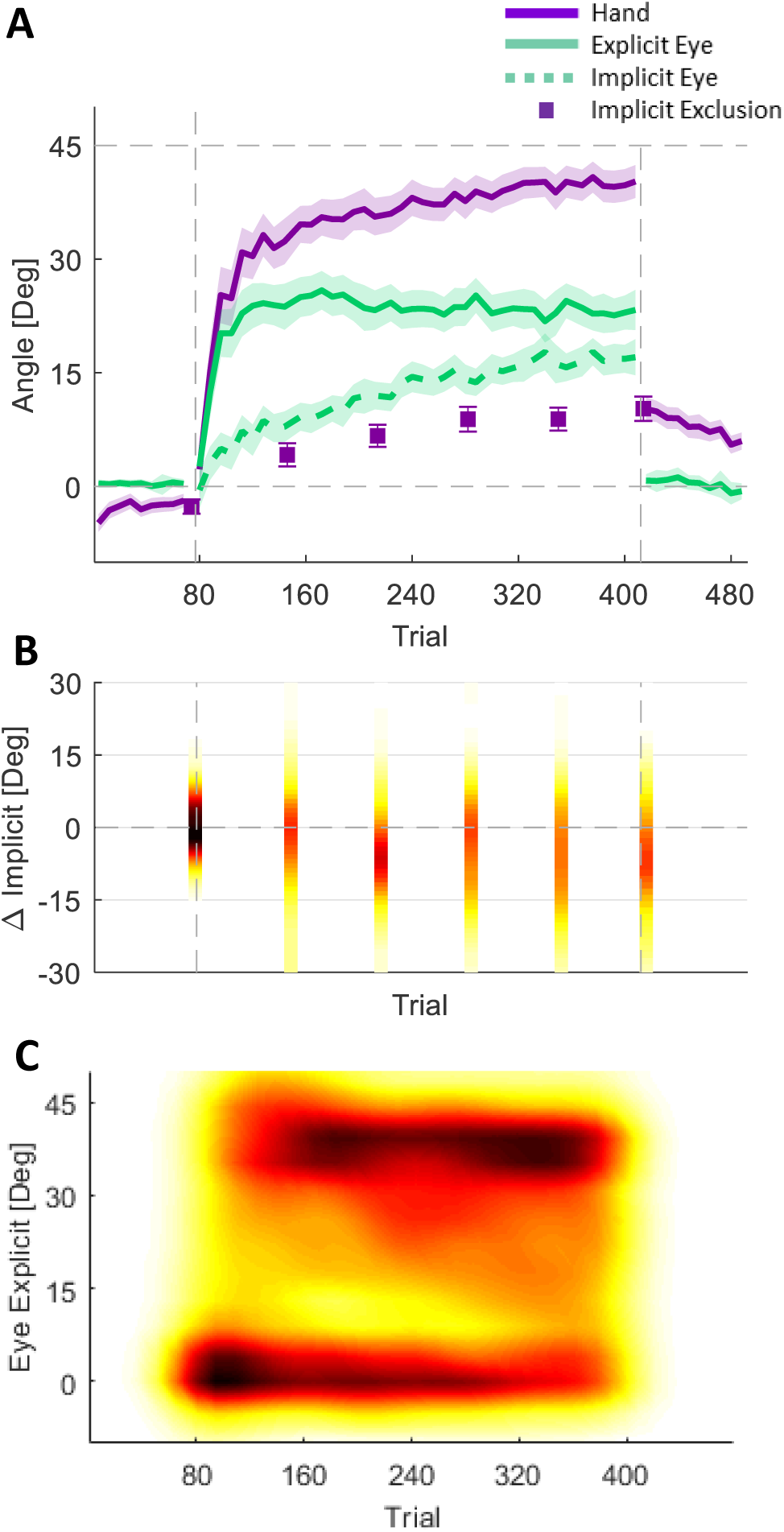
Experiment 2. (**A**) Averaged learning curves across the entire group. Error bars in the Implicit Exclusion and the shaded area represent the SEM. (**B**) Distribution of the differences between the two implicit measures. (**C**) Distribution of the Explicit Eye.

The learning curve of the different subjects exemplify fundamental differences between subjects. First, for some subjects, the two measures of implicit correspond while, for others, the Implicit Eye was higher than the Implicit Exclusion (Figure 4B). Second, within subjects with corresponding measures of implicit: for some gaze stayed locked on the target, reflecting an Explicit Eye adaptation of 0° (these subjects also had a very slow rise in Hand-Target Difference which reaches no more than 30°); while for others, gaze shifted rapidly to the hand-target, reflecting nearly full explicit knowledge (Figure 4C) and it was consistent with a quick rise in the Hand-Target Difference.

Having noticed this pattern in the data, we clustered the subjects accordingly (Figure 5). We used a two-step approach. In the first step, we clustered subjects using FCM according to the difference between the two measures of Implicit. The cluster validity index (Zhang et al., 2008) values suggested two clusters, As expected, for one cluster Implicit Eye and Implicit Exclusion matched, while for the other cluster Implicit Exclusion was greater than Implicit Eye. We called these the Matched-Implicit and No-Match clusters. In the second clustering step, we further clustered the Matched-Implicit cluster according to the Explicit Eye measured through the entire rotation block. We used PPCA to reduce the dimensionality of the data. We ran FCM clustering over the first three components which captured 93% of the overall variance. The cluster validity values again suggested two clusters, while when applying the same clustering approach to the No-Match group they did not reveal any clustering.

**Figure 5.**
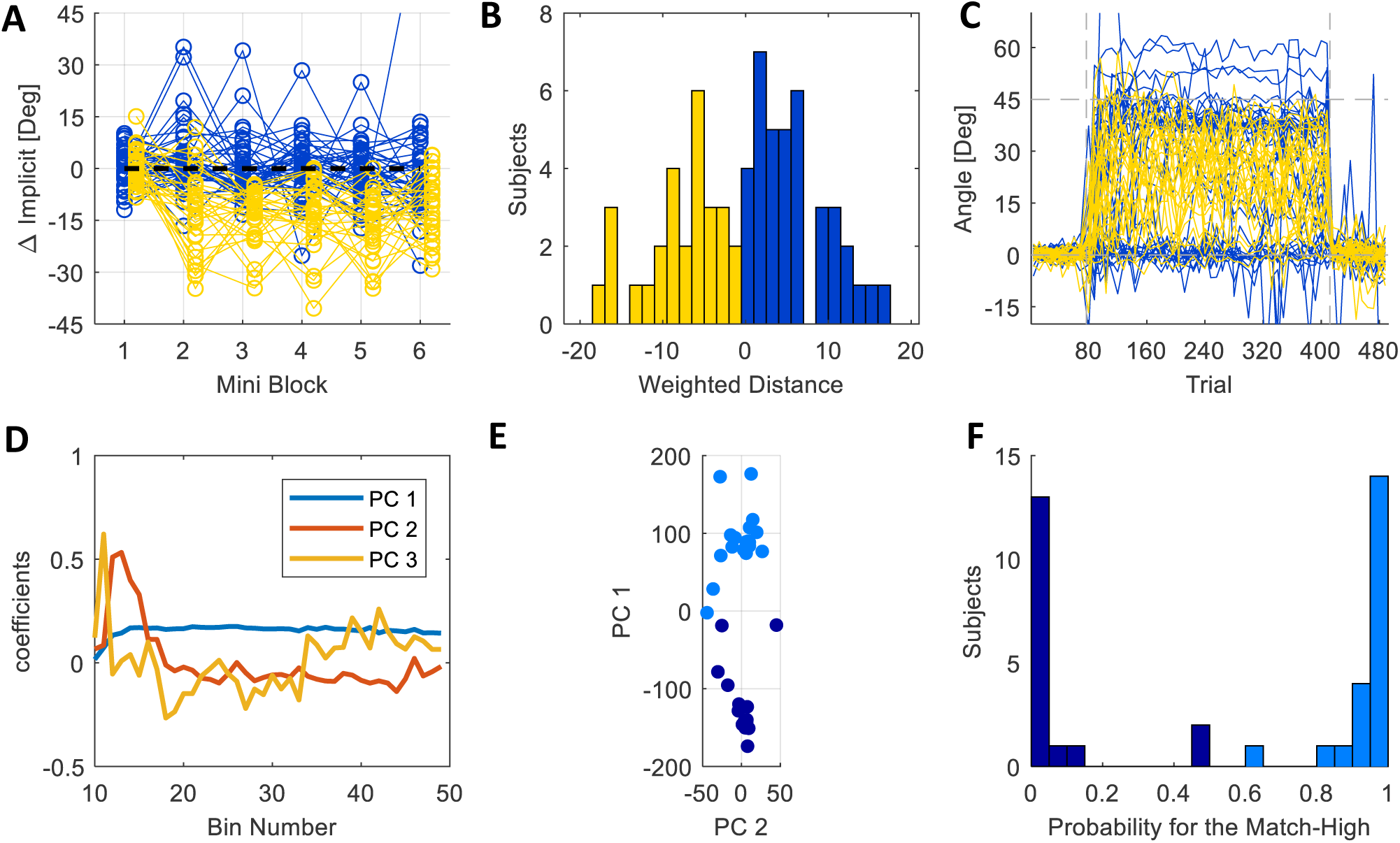
Experiment 2: Clustering steps. (**A-C**) Step 1. Blue lines are subjects who belong to the cluster in which the Implicit measures matches and yellow line to the cluster the implicit measures don’t match: (**A**) The difference between the two implicit measures for the six Mini Exclusion Blocks. Each line depicts an individual subject. (**B**) Histogram of the weighted averaged Implicit difference. (**C**) Explicit Eye for all subjects. (**D-F**) Step 2. (**D**) 3 principal components, on which the data is clustered, are shown. (**E**) The 1st and the 2nd PCs. Dots represent individual subject. Light blue are subjects with high Explicit Eye and dark blue are subject with low Explicit Eye. (**F**) Histogram of the probability of subjects to belong to the High Explicit Eye cluster.

The two-step clustering method categorized subjects into three clusters. Two of them are derived from the Matched-Implicit cluster of the first step, and, for them, the Explicit Eye faithfully reflects explicit knowledge. From those two, we called the one that had more explicit adaptation Match-High (n=21). The Hand-Target Difference of these subjects rose quickly to 31.8 ± 6.6° during the initial rise, and then to 45.4 ± 2.2° at the late early rise, and at the end of adaptation stand on 48.7 ± 2.4° (Figure 6A). This group of subjects counteract the rotation fully and quickly using an explicit strategy. This can be seen in the Explicit Eye which rose quickly to 31.0 ± 6.2°, later to 41.6 ± 2.7°, and by the end of adaptation was slightly reduced to 38.8 ± 2.3° (Figure 6B). Their implicit learning, measured by Implicit Eye, was 10.6 ± 2.8° at the end of the rotation block (Figure 6C). For this group, Explicit Eye and Implicit Eye matched Explicit Exclusion and Implicit Exclusion (differences of 0.5 ± 1.1° and −0.3 ± 1.6°, respectively).

**Figure 6.**
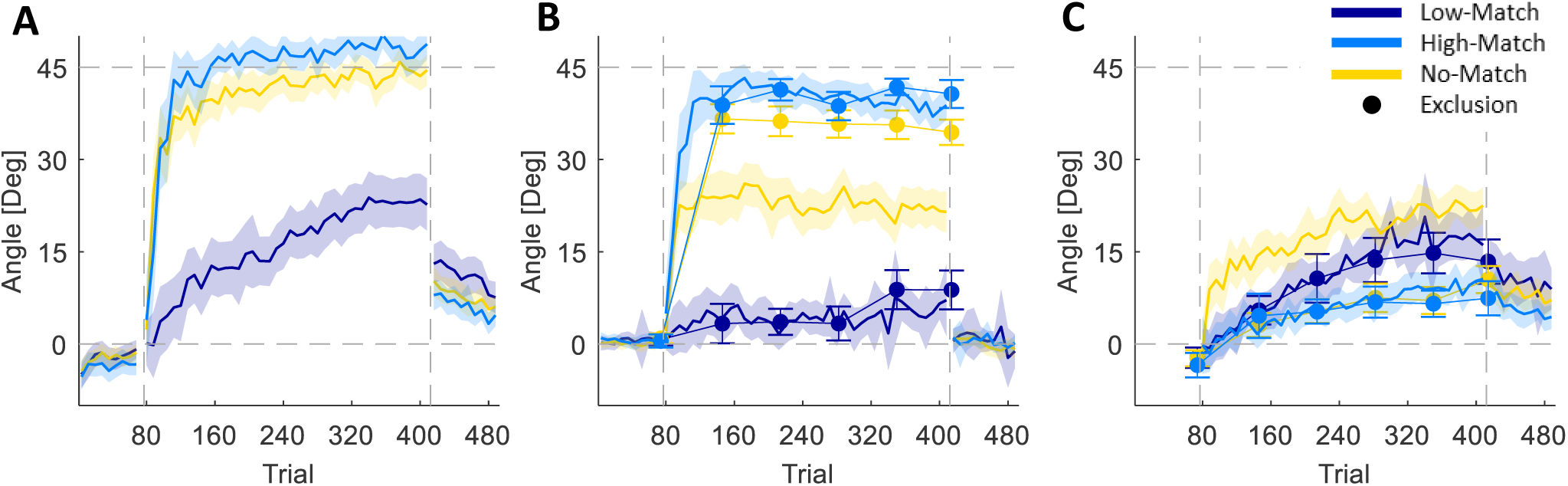
Experiment 2: Learning curves per cluster. (**A**) Hand Target Difference for the Match-Low, Match-High and No-Match clusters. (**B**) Explicit Eye (lines) and Explicit Exclusion (circles) for the three clusters. (**C**) Implicit Eye (lines) and Implicit Exclusion (circles) for the three clusters. Error bars around the Implicit Exclusion and the shaded area represent the SEM.

In contrast, the other matching cluster had very little explicit adaptation. We called it Match-Low (n=17). For these subjects, the difference between Explicit Eye and Explicit Exclusion was also very small (−0.1 ± 1.9°), as was the case for Implicit Eye and Implicit Exclusion (−0.5 ± 2.4°. The lack of explicit strategy is reflected in the very low Explicit Eye which was 3.0 ± 4.2° in the initial rise, 5.8 ± 4.0° at the late early rise, and reached to 7.1 ± 5.9° at the end of adaptation (Figure 6B). Since this cluster showed no explicit strategy, they adapt to the rotation only implicitly, and thus have very slow adaptation (Figure 6A). The average Hand-Target Difference changed from 3.6 ± 5.6° at the initial rise to 12.7 ± 4.7° at the late early rise and reached 22.7 ± 4.3° at the end of the rotation block. Hence, the Match-High and Match-Low groups had very different Hand-Target Difference and Explicit Eye. The Implicit Eye of the Match-Low group reached 16.1 ± 6.5° by the end of the rotation block, 5.5 ± 7.1° higher than that of the Match-High group.

The No-Match (n=28; which was the second cluster in step 1) was characterize by a lack of correspondence between Implicit Eye and Implicit Exclusion (Figure 6C). Accordingly, this cluster adapts to the rotation using mostly explicit strategy (Figure 6B). However, the Explicit Eye captures only part of the explicit learning measured with Explicit Exclusion. The difference between them, across subjects, is 13.5 ± 1.5° by the end of the rotation block. The fact that explicit adaptation does exist in these subjects is supported by the rapid rise of the Hand-Target Difference: 33.5 ± 5.0° in the initial rise, 39.8 ± 2.8° at the late early rise, and 44.5 ± 2.5° at the end of adaptation. This performance is similar to that of the Match-High group, with differences of −1.7 ± 8.2°, 5.6 ± 3.6° and 4.2 ± 3.5° at the initial rise, late early rise and at the end of adaptation. In contrast, the Explicit Eye of this No-Match group was very different from that of the Match-High. It changed from 22.6 ± 4.1° in the initial rise to 24.1 ± 3.6° at the late early rise and to 21.5 ± 3.3° by the end of adaptation. Both measures of implicit of the No-Match cluster rose slowly; however, the Implicit Eye was, obviously, much higher than Implicit Exclusion, by 13.3 ± 1.7° by the end of the rotation block.

Following previous studies that suggest that explicit learning requires longer RTs (Benson et al., 2011 e.g.; Leow et al., 2017; McDougle and Taylor, 2019), we looked for differences in the RT between our different explicit learning groups (Figure 7A). The Match-High group showed the highest RTs. Their RTs decreased by 758 ± 370ms from initial adaptation to the end of adaptation, in line with the decrease in their Explicit Eye over the adaptation period. The RTs of the Match-Low group, who had almost no explicit adaptation, was much smaller on average. Though the Explicit Eye of the Match-Low increased during adaptation, their RTs decreased by 450 ± 341ms. The RTs of the No-Match group started as low as the Match-Low, over adaptation it dropped only by 272 ± 228ms. Interestingly, the Match-High group had longer RTs in the baseline and the washout blocks as well. The RTs of this group were different from those of the Low-match and the No-Match by 249 ± 181ms and 404 ± 152ms, respectively. This suggests that behavior in the clusters may be different even before adaptation begins.

**Figure 7.**
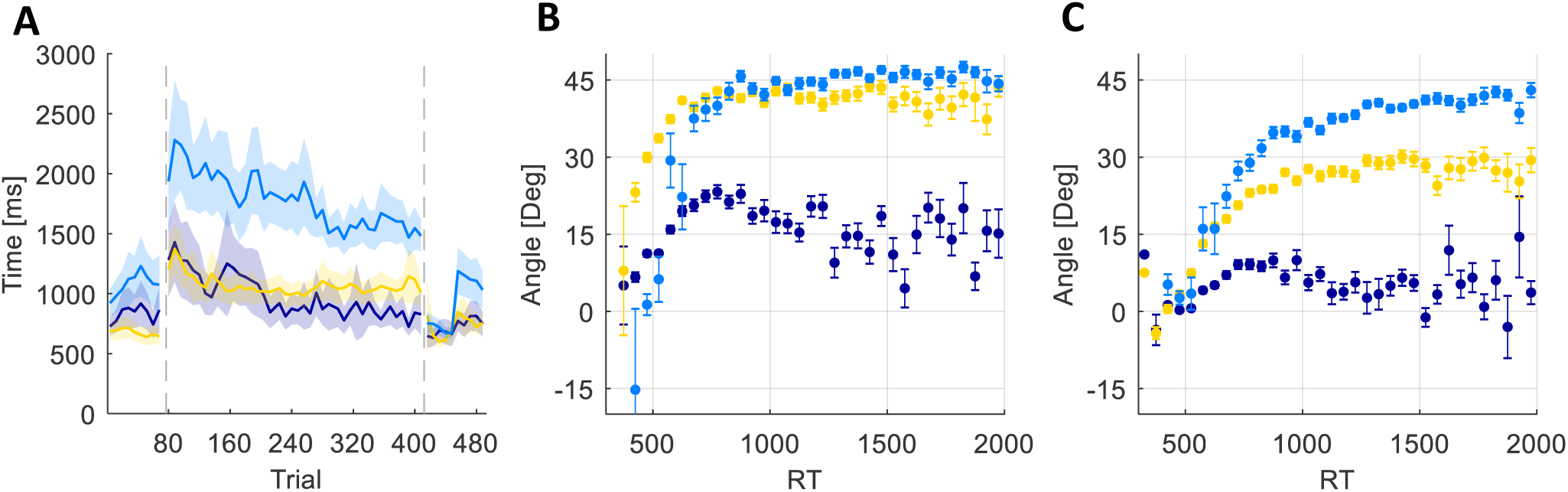
Experiment 2: Reaction times. (**A**) Averaged reaction time for each cluster. The shaded area represents the SEM. (**B**) Cluster means of Hand Target Differences and (**C**) cluster means of Explicit Eye as a function of RT (binned in 25ms bins). Each dot is a bin and error bars represent SEM of each bin

Figure 7B and C show the relationship between RT and Hand-Target Difference and between RT and Explicit Eye. For RTs higher than 400ms, larger RTs were associated with larger Hand-Target Differences and with more Explicit Eye. In the Match-Low group this relationship saturated around 700ms. While the general picture supports the idea that Explicit Eye reflects aspects of explicit knowledge, this idea would also imply that averaged RTs in the No-Match group should be closer to those of the Match-High rather than to those of the Match-Low, and this is contradicted by Figure 7A. For this group the Explicit Eye reflects only one component of explicit learning, the same one that is correlated to the RTs, while there is another explicit component which is not related to either eye movements or RT.

## Discussion

In this study, we explored the extent to which explicit components of visuomotor adaptation are captured by eye movements. We did this by comparing eye movements to two accepted measures of explicit learning – verbal report and the exclusion test. Our experiments showed that eye movements have a stable pattern: after target appearance the eyes saccade from the origin to the target, and then, before movement onset, the eyes saccade again in the direction towards which the subject will aim. We believe that these eye movements provide a measure of explicit adaptation (we called it Explicit Eye); however, this measure only reflects part of the explicit adaptation. Our first experiment showed that when subjects report their intended direction, Explicit Eye and the other two measures (verbal report and exclusion) all matched. In contrast, when subjects did not report, Explicit Eye only reflected part of the explicit adaptation indicated by the exclusion. The fact that the two were correlated suggested that Explicit Eye might be reflecting components of explicit adaptation (Figure 3F). In our second experiment, we tried to explore more fully the time course of the separation of Explicit Eye from explicit shown by exclusion. We found that the two diverge early in adaptation. In analyzing the data of the second experiment, we found three groups of subjects. The first group adapted fully to the rotation and had eye movements consistent with performance in exclusion trials (Match-High); the second group also adapted fully but had less Explicit Eye than would be expected from exclusion trials (No-Match); the third group only adapted partially and had eye movements consistent with lack of explicit adaptation in the exclusion trials (Match-Low). The learning curves of this last group were similar to those reported in paradigms were subjects had only implicit adaptation (Kim et al., 2018, 2019; Morehead et al., 2017).

Smith et al. (2006) proposed a two-state model of motor adaptation that is still the most widely used model in the field. It has been nicely mapped onto explicit (fast) and implicit (slow) components of adaptation (McDougle et al., 2015). However, there have been suggestions that there are more than two states in the adaptation process and that there may be multiple explicit and implicit components, potentially with different time constants. Forano and Franklin (2019) showed that dual-adaptation can be best explained by models with two fast components, and McDougle and Taylor (2019) showed two explicit strategies in visuomotor rotation: caching and mental rotation. Presumably, in our study, the group for which Explicit Eye explained only part of the explicit learning (No-Match group) used multiple explicit strategies, while the group where measures of explicit learning matched used one (Match-High) or, perhaps, none (Match-Low).

The question arises whether the components of the explicit adaptation that are reflected in Explicit Eye map to the explicit strategies identified by McDougle and Taylor (2019). In that study, the key difference in the strategies was that one strategy introduced a correlation between rotation and reaction time while the other did not. Consequently, we examined reaction times in the different groups. We found that the group with the single explicit strategy (captured by gaze; Match-High) had very long reaction times relative to the other groups. Interestingly, these subjects had longer reaction times in the baseline phase as well, suggesting that they were more carefully and explicitly controlled movers even during normal movement. The reaction times of the No-Match group were much lower. That is, the No-Match group achieved explicit adaptation comparable to the Match-High group, but their explicit adaptation did not require preparation time. The Match-Low subjects, who on had implicit adaptation, had the fastest reaction times. Taken together, these results suggest that the explicit components reflected in Explicit Eye are the same components that drive longer reaction times. McDougle and Taylor (2019) identified this as the process of mental rotation and contrasted it with the low reaction time mechanism of caching.

Links between intended direction of movement and eye movements have been foreshadowed (Rand and Rentsch, 2016; Rentsch and Rand, 2014) and demonstrated explicitly (de Brouwer et al., 2018). Our study supports these earlier findings, although there are some technical issues that deserve consideration. First, we followed the Rand and Rentsch (2016) study in using only end-point feedback rather than continuous presentation of the cursor. This simplified the eye movements and allowed us to determine that the fixations immediately prior to movement initiation provided the most reliable estimate of explicit adaptation. The specific timing at which eye movements are considered leads to a second issue. Despite a general agreement between our study and de Brouwer et al. (2018) that eye movements reflect explicit adaptation, there is an important difference in the findings that relates to the timing at which eye movements are considered. De Brouwer et al. evaluate the fixation closest to the rotation angle. Our data suggest that this measure overestimates the explicit adaptation (as measured by the exclusion test). The last fixation before movement onset was a more stable measure and is consistent with earlier results on the specific timing with which eye movements predict hand movements (Ariff et al., 2002). This measure also allowed an unbiased quantification of explicit adaptation in subjects with very little of it, a key aspect in identifying out Match-Low group. This difference in the measures may be one reason why de Brouwer et al. (2018) did not see the three different groups of subjects that we did.

### Conclusions

This study provides additional support to earlier findings that eye movements reflect explicit strategy in visuomotor adaptation. However, it also highlights the existence of different explicit components. It seems that some components of explicit adaptation are not reflected in the eye movements. The component that is reflect in the eye movements is correlated with reaction time and may be the same component identified by McDougle and Taylor (2019) as mental rotation. While eye movements may not be a perfect measure of explicit adaptation, they could be used to capture this component on a trial-by-trial basis without influencing the adaptation.

## Acknowledgements

We would like to thank Ilan Dinstein and Ayelet Arazi for their help with the eye tracking. This work was partially supported by DFG grant TI-239/16-1. Shlomi Haar is supported by the Royal Society – Kohn International Fellowship (NF170650)

## Statistical Appendix

The next tables present the statistical analysis. The *μ*, *σ*, Δ and effect size computed on the original Data. The CI and probability to reject null computed on the sampled distribution.

Effect size is calculated either by 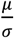 or by 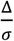.

### Experiment 1

#### Initial rise in Hand-Target Difference

Initial rise in Hand-Target Difference:
| group | $\mu$ | $\sigma$ | 2.5% | 97.5% | Prob to reject null ($\Delta > 0$) | Effect size |
| --- | --- | --- | --- | --- | --- | --- |
| NR | 24.7 | 20.8 | 8.2 | 41.3 | 0.18% | 1.2 |
| R | 32.9 | 22.5 | 19.1 | 43.6 | 0% | 1.5 |

#### Differences between groups

Differences between groups:
| $\Delta$ | 2.5% | 97.5% | Prob to reject null | | Effect size |
| --- | --- | --- | --- | --- | --- |
| | | | $\Delta > 0$ | $\Delta = 0$ | |
| 8.2 | -12.7 | 28.8 | 21% | 73% | 1.7 |

#### Late early rise in Hand-Target Difference

Late early rise in Hand-Target Difference:
| group | $\mu$ | $\sigma$ | 2.5% | 97.5% | Prob to reject null ($\Delta > 0$) | Effect size |
| --- | --- | --- | --- | --- | --- | --- |
| NR 1<sup>st</sup> | 33.9 | 13.7 | 22.7 | 43.9 | 0% | 2.5 |
| R 1<sup>st</sup> | 46.5 | 5.6 | 40.9 | 52.1 | 0% | 8.2 |

#### Differences between groups

Differences between groups:
| $\Delta$ | 2.5% | 97.5% | Prob to reject null | | Effect size |
| --- | --- | --- | --- | --- | --- |
| | | | $\Delta > 0$ | $\Delta = 0$ | |
| 12.6 | 0.9 | 25.2 | 2% | 73% | 4.1 |

#### End of adaptation in the Hand-Target Difference

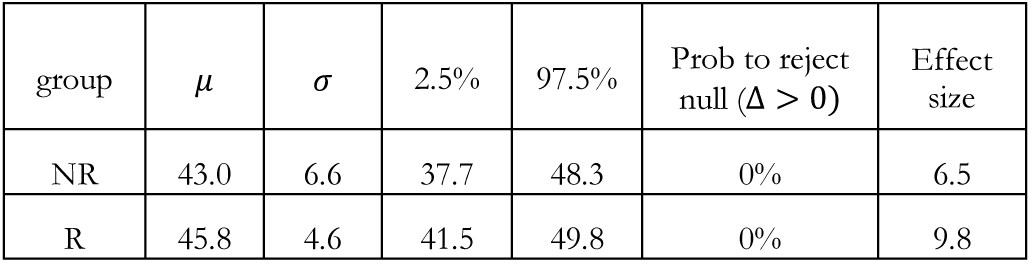

End of adaptation in the Hand-Target Difference:
| group | $\mu$ | $\sigma$ | 2.5% | 97.5% | Prob to reject null ($\Delta > 0$) | Effect size |
| --- | --- | --- | --- | --- | --- | --- |
| NR | 43.0 | 6.6 | 37.7 | 48.3 | 0% | 6.5 |
| R | 45.8 | 4.6 | 41.5 | 49.8 | 0% | 9.8 |

#### Differences between groups

Differences between groups:
| $\Delta$ | 2.5% | 97.5% | Prob to reject null | | Effect size |
| --- | --- | --- | --- | --- | --- |
| | | | $\Delta > 0$ | $\Delta = 0$ | |
| 2.8 | -3.9 | 9.4 | 21% | 27% | 1.2 |

#### Initial rise in Explicit Eye

Initial rise in Explicit Eye:
| group | $\mu$ | $\sigma$ | 2.5% | 97.5% | Prob to reject null ($\Delta > 0$) | Effect size |
| --- | --- | --- | --- | --- | --- | --- |
| NR | 17.7 | 17.4 | 5.6 | 30.1 | 0.23% | 1.0 |
| R | 33.6 | 22.3 | 21.1 | 43.9 | 0% | 1.5 |

#### Differences between groups

Differences between groups:
| $\Delta$ | 2.5% | 97.5% | Prob to reject null | | Effect size |
| --- | --- | --- | --- | --- | --- |
| | | | $\Delta > 0$ | $\Delta = 0$ | |
| 15.8 | -1.4 | 32.0 | 4% | 91% | 3.4 |

#### Late early rise in Explicit Eye

**Late early rise in Explicit Eye:**
| group | $\mu$ | $\sigma$ | 2.5% | 97.5% | Prob to reject null ($\Delta > 0$) | Effect size |
| --- | --- | --- | --- | --- | --- | --- |
| NR | 24.7 | 14.8 | 13.2 | 35.6 | 0% | 1.6 |
| R | 44.2 | 5.1 | 41.0 | 47.4 | 0% | 8.6 |

#### Differences between groups

**Differences between groups:**
| $\Delta$ | 2.5% | 97.5% | Prob to reject null | | Effect size |
| --- | --- | --- | --- | --- | --- |
| | | | $\Delta > 0$ | $\Delta = 0$ | |
| 19.5 | 7.9 | 31.4 | 0.05% | 99% | 6.3 |

#### End of adaptation in Explicit Eye

**End of adaptation in Explicit Eye:**
| group | $\mu$ | $\sigma$ | 2.5% | 97.5% | Prob to reject null ($\Delta > 0$) | Effect size |
| --- | --- | --- | --- | --- | --- | --- |
| NR | 18.9 | 14.4 | 11.1 | 27.3 | 0% | 1.3 |
| R | 36.8 | 9.6 | 31.7 | 41.7 | 0% | 3.8 |

#### Differences between groups

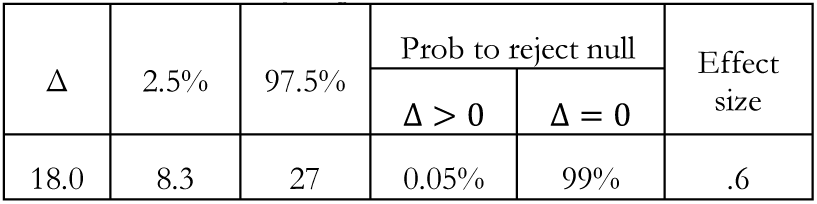

#### Initial rise in Explicit Report

**Initial rise in Explicit Report:**
| | $\mu$ | $\sigma$ | 2.5% | 97.5% | Prob to reject null ($\Delta > 0$) | Effect size |
| --- | --- | --- | --- | --- | --- | --- |
| R | 32.4 | 23.1 | 18.7 | 43.9 | 0% | 1.4 |

#### Late early rise in Explicit Report

**Late early rise in Explicit Report:**
| | $\mu$ | $\sigma$ | 2.5% | 97.5% | Prob to reject null ($\Delta > 0$) | Effect size |
| --- | --- | --- | --- | --- | --- | --- |
| R | 44.3 | 7.1 | 40.8 | 48.6 | 0% | 6.2 |

#### End of adaptation in Explicit Report

**End of adaptation in Explicit Report:**
| | $\mu$ | $\sigma$ | 2.5% | 97.5% | Prob to reject null ($\Delta > 0$) | Effect size |
| --- | --- | --- | --- | --- | --- | --- |
| R | 37.7 | 9.1 | 33.5 | 41.5 | 0% | 4.1 |

#### Report explicit - eye explicit within a group

**Report explicit - eye explicit within a group:**
| | $\Delta$ | 2.5% | 97.5% | Prob to reject null ($\Delta=0$) | Effect size |
| --- | --- | --- | --- | --- | --- |
| R | -0.02 | -1.6 | 1.5 | 0% | 0 |

#### Report implicit - eye implicit within a group

**Report implicit - eye implicit within a group:**
| | $\Delta$ | 2.5% | 97.5% | Prob to reject null ($\Delta=0$) | Effect size |
| --- | --- | --- | --- | --- | --- |
| R | -0.3 | -1.9 | 1.4 | 0% | -0.3 |

#### Eye Implicit - After effect within a group

**Eye Implicit - After effect within a group:**
| Group | $\Delta$ | 2.5% | 97.5% | Prob to reject null ($\Delta=0$) | Effect size |
| --- | --- | --- | --- | --- | --- |
| NR | 19.6 | 10.8 | 28.4 | 100% | 1.7 |
| R | 3.4 | -2.2 | 9.3 | 29% | 0.5 |

#### Differences between groups

**Differences between groups:**
| $\Delta$ | 2.5% | 97.5% | Prob to reject null (R>NR) | Effect size |
| --- | --- | --- | --- | --- |
| -16.2 | -26.5 | -5.8 | 0% | -1.8 |

#### after effect Vs implicit eye: correlation

| Group | Mean correlation | 2.5% | 97.5% | Prob to reject null ($r > 0.2$) |
| --- | --- | --- | --- | --- |
| NR | 0.67 | 0.36 | 0.79 | 0.1% |
| R | 0.75 | 0.49 | 0.83 | 0% |

### Experiment 2

#### Implicit Eye – Catch

Experiment 2: Implicit Eye – Catch:
| cluster | $\Delta$ | $\sigma$ | 2.5% | 97.5% | Prob to reject null ($\Delta = 0$) | Effect size |
| --- | --- | --- | --- | --- | --- | --- |
| all | 5.4 | 8.1 | 2.7 | 8.1 | 62% | 0.6 |
| Match Low | -0.5 | 4.5 | -5.6 | 4.0 | 5% | -0.1 |
| No Match | 13.3 | 4.7 | 10.0 | 16.7 | 100% | 2.8 |
| Match High | -0.3 | 4.3 | -3.5 | 2.8 | 0.3% | -0.06 |

#### Explicit Eye – Catch

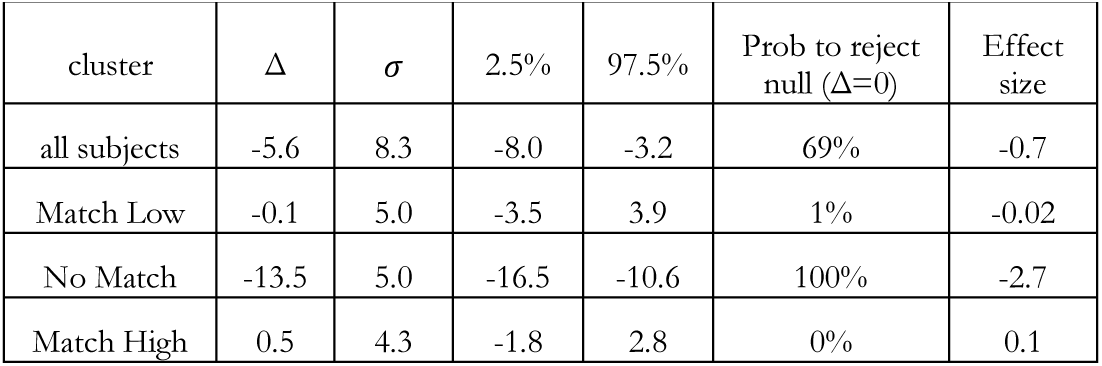

#### Initial rise in Hand-Target Difference

Initial rise in Hand-Target Difference:
| group | $\mu$ | $\sigma$ | 2.5% | 97.5% | Prob to reject null ($\Delta > 0$) | Effect size |
| --- | --- | --- | --- | --- | --- | --- |
| all | 25.2 | 21.0 | 17.9 | 32.4 | 0% | 1.2 |
| Match Low | 3.6 | 13.7 | -7.8 | 14.7 | 25% | 0.3 |
| No Match | 33.5 | 16.0 | 22.8 | 42.5 | 0% | 2.1 |
| Match High | 31.8 | 20.0 | 18.0 | 44.4 | 0% | 1.6 |

#### Differences between clusters

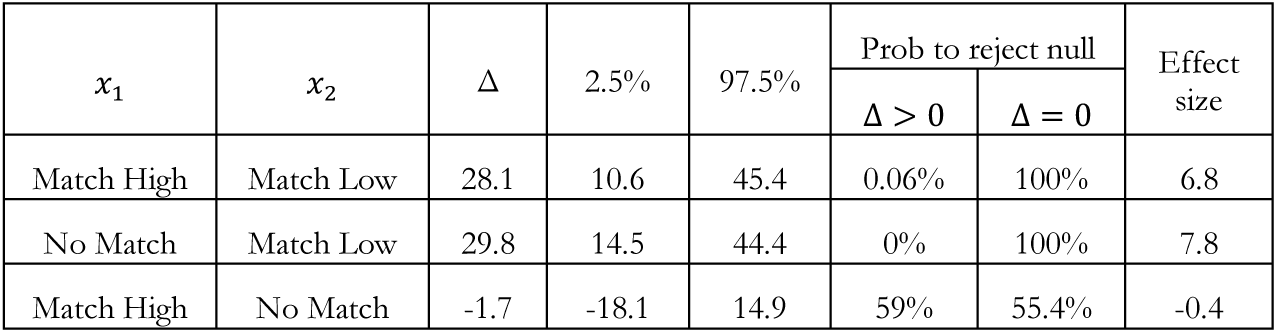

Differences between clusters:
| $x_1$ | $x_2$ | $\Delta$ | 2.5% | 97.5% | Prob to reject null | | Effect size |
| --- | --- | --- | --- | --- | --- | --- | --- |
| | | | | | $\Delta > 0$ | $\Delta = 0$ | |
| Match High | Match Low | 28.1 | 10.6 | 45.4 | 0.06% | 100% | 6.8 |
| No Match | Match Low | 29.8 | 14.5 | 44.4 | 0% | 100% | 7.8 |
| Match High | No Match | -1.7 | -18.1 | 14.9 | 59% | 55.4% | -0.4 |

#### Late early rise in Hand-Target Difference

Late early rise in Hand-Target Difference:
| group | $\mu$ | $\sigma$ | 2.5% | 97.5% | Prob to reject null ($\Delta > 0$) | Effect size |
| --- | --- | --- | --- | --- | --- | --- |
| all | 34.6 | 26.6 | 29.7 | 32.3 | 0% | 2.1 |
| Match Low | 12.7 | 14.5 | 4.3 | 22.2 | 0.1% | 0.9 |
| No Match | 39.8 | 10.2 | 34.0 | 45.1 | 0% | 3.9 |
| Match High | 45.4 | 4.7 | 41.0 | 49.7 | 0% | 9.6 |

#### Differences between clusters

Differences between clusters:
| $x_1$ | $x_2$ | $\Delta$ | 2.5% | 97.5% | Prob to reject null | | Effect size |
| --- | --- | --- | --- | --- | --- | --- | --- |
| | | | | | $\Delta > 0$ | $\Delta = 0$ | |
| Match High | Match Low | 32.7 | 22.3 | 42.3 | 0% | 100% | 10.9 |
| No Match | Match Low | 27.0 | 15.7 | 37.2 | 0% | 100% | 7.9 |
| Match High | No Match | 5.6 | -1.3 | 12.7 | 6% | 57% | 2.0 |

#### End of adaptation in Hand-Target Difference

**End of adaptation in Hand-Target Difference:**
| group | $\mu$ | $\sigma$ | 2.5% | 97.5% | Prob to reject null ($\Delta > 0$) | Effect size |
| --- | --- | --- | --- | --- | --- | --- |
| all | 40.2 | 14.0 | 35.9 | 44.4 | 0% | 2.99 |
| Match Low | 22.7 | 14.4 | 14.3 | 31.0 | 0% | 1.5 |
| No Match | 44.5 | 7.4 | 39.4 | 49.3 | 0% | 6.0 |
| Match High | 48.7 | 5.6 | 43.8 | 53.5 | 0% | 8.7 |

#### Differences between clusters

Differences between clusters:
| $x_1$ | $x_2$ | $\Delta$ | 2.5% | 97.5% | Prob to reject null | | Effect size |
| --- | --- | --- | --- | --- | --- | --- | --- |
| | | | | | $\Delta > 0$ | $\Delta = 0$ | |
| Match High | Match Low | 26.1 | 16.3 | 35.8 | 0% | 100% | 8.5 |
| No Match | Match Low | 21.8 | 12.0 | 31.4 | 0% | 100% | 6.9 |
| Match High | No Match | 4.2 | -2.6 | 11.2 | 11.2% | 42% | 1.6 |

#### Initial rise in Explicit Eye

**Initial rise in Explicit Eye:**
| group | $\mu$ | $\sigma$ | 2.5% | 97.5% | Prob to reject null ($\Delta > 0$) | Effect size |
| --- | --- | --- | --- | --- | --- | --- |
| all | 20.2 | 18.7 | 14.2 | 26.5 | 0% | 1.1 |
| Match Low | 3.0 | 9.4 | -4.7 | 11.9 | 23% | 0.3 |
| No Match | 22.6 | 148 | 14.6 | 30.4 | 0% | 1.5 |
| Match High | 31.0 | 19.8 | 19.0 | 43.0 | 0% | 1.6 |

#### Differences between clusters

Differences between clusters:
| $x_1$ | $x_2$ | $\Delta$ | 2.5% | 97.5% | Prob to reject null | | Effect size |
| --- | --- | --- | --- | --- | --- | --- | --- |
| | | | | | $\Delta > 0$ | $\Delta = 0$ | |
| Match High | Match Low | 28.1 | 13.0 | 42.4 | 0% | 100% | 7.2 |
| No Match | Match Low | 19.6 | 7.7 | 30.5 | 0.1% | 99% | 5.5 |
| Match High | No Match | 8.5 | -6.0 | 23.0 | 12% | 73% | 2.1 |

#### Late early rise in Explicit Eye

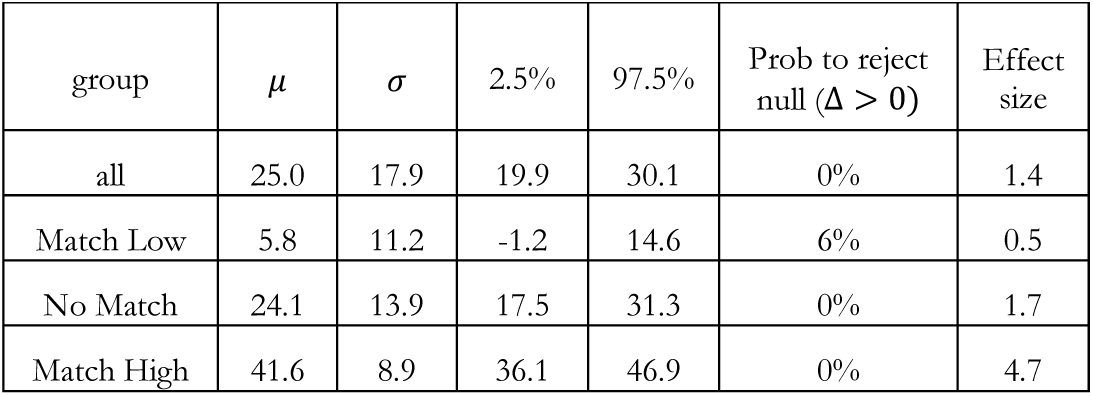

#### Differences between clusters

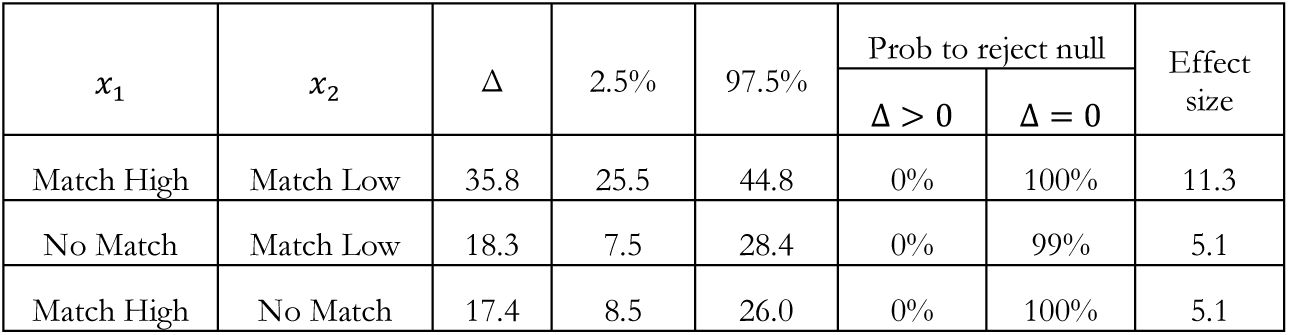

Differences between clusters:
| $x_1$ | $x_2$ | $\Delta$ | 2.5% | 97.5% | Prob to reject null | | Effect size |
| --- | --- | --- | --- | --- | --- | --- | --- |
| | | | | | $\Delta > 0$ | $\Delta = 0$ | |
| Match High | Match Low | 35.8 | 25.5 | 44.8 | 0% | 100% | 11.3 |
| No Match | Match Low | 18.3 | 7.5 | 28.4 | 0% | 99% | 5.1 |
| Match High | No Match | 17.4 | 8.5 | 26.0 | 0% | 100% | 5.1 |

#### End of adaptation in Explicit Eye

**End of adaptation in Explicit Eye:**
| group | $\mu$ | $\sigma$ | 2.5% | 97.5% | Prob to reject null ($\Delta > 0$) | Effect size |
| --- | --- | --- | --- | --- | --- | --- |
| all | 23.3 | 16.8 | 18.3 | 28.6 | 0% | 1.4 |
| Match Low | 7.1 | 14.1 | -2.0 | 21.0 | 6% | 0.5 |
| No Match | 21.5 | 12.0 | 14.8 | 28.1 | 0% | 1.8 |
| Match High | 38.8 | 9.0 | 34.3 | 43.3 | 0% | 4.3 |

#### Differences between clusters

Differences between clusters:
| $x_1$ | $x_2$ | $\Delta$ | 2.5% | 97.5% | Prob to reject null | | Effect size |
| --- | --- | --- | --- | --- | --- | --- | --- |
| | | | | | $\Delta > 0$ | $\Delta = 0$ | |
| Match High | Match Low | 31.7 | 17.1 | 42.0 | 0% | 100% | 9.4 |
| No Match | Match Low | 14.3 | -0.7 | 25.7 | 3% | 92% | 4.0 |
| Match High | No Match | 17.3 | 9.3 | 25.2 | 0% | 100% | 5.3 |

#### End of adaptation in Implicit Eye

End of adaptation in Implicit Eye:
| group | $\mu$ | $\sigma$ | 2.5% | 97.5% | Prob to reject null ($\Delta > 0$) | Effect size |
| --- | --- | --- | --- | --- | --- | --- |
| all | 17.0 | 12.4 | 12.2 | 21.4 | 0% | 1.4 |
| Match Low | 16.1 | 16.7 | 1.2 | 27.0 | 2% | 1.0 |
| No Match | 22.5 | 9.5 | 16.7 | 28.1 | 0% | 2.3 |
| Match High | 10.6 | 8.3 | 5.0 | 16.2 | 0% | 1.3 |

#### Differences between clusters

Differences between clusters:
| $x_1$ | $x_2$ | $\Delta$ | 2.5% | 97.5% | Prob to reject null | | Effect size |
| --- | --- | --- | --- | --- | --- | --- | --- |
| | | | | | $\Delta < 0$ | $\Delta = 0$ | |
| Match High | Match Low | -5.5 | -18.0 | 10.1 | 20% | 65% | 1.0 |
| No Match | Match Low | 6.4 | -6.1 | 22.4 | 83% | 59% | 2.4 |
| Match High | No Match | -11.9 | -19.8 | -3.7 | 0% | 95% | 1.3 |

#### Decrease in RT

Decrease in RT:
| group | $\mu$ | $\sigma$ | 2.5% | 97.5% | Prob to reject null ($\Delta > 0$) | Effect size |
| --- | --- | --- | --- | --- | --- | --- |
| Match Low | 450 | 820 | -189 | 1182 | 8% | 0.5 |
| No Match | 272 | 497 | -162 | 733 | 11% | 0.5 |
| Match High | 758 | 1066 | 73 | 1537 | 1% | 0.7 |

#### RT in baseline

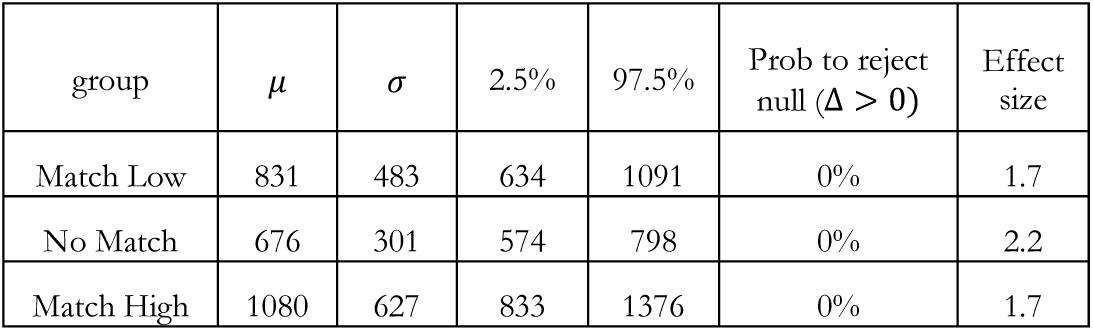

RT in baseline:
| group | $\mu$ | $\sigma$ | 2.5% | 97.5% | Prob to reject null ($\Delta > 0$) | Effect size |
| --- | --- | --- | --- | --- | --- | --- |
| Match Low | 831 | 483 | 634 | 1091 | 0% | 1.7 |
| No Match | 676 | 301 | 574 | 798 | 0% | 2.2 |
| Match High | 1080 | 627 | 833 | 1376 | 0% | 1.7 |

#### Differences between clusters

Differences between clusters:
| $x_1$ | $x_2$ | $\Delta$ | 2.5% | 97.5% | Prob to reject null | | Effect size |
| --- | --- | --- | --- | --- | --- | --- | --- |
| | | | | | $\Delta < 0$ | $\Delta > 0$ | |
| Match High | Match Low | 249 | -104 | 610 | 8% | 91% | 1.7 |
| No Match | Match Low | -155 | -433 | 79 | 89% | 10% | 2.2 |
| Match High | No Match | 404 | 130 | 725 | 0.1% | 100% | 1.7 |

## References

Amrhein, V., Greenland, S., and McShane, B. (2019). Scientists rise up against statistical significance. Nature 567, 305–307.

Ariff, G., Donchin, O., Nanayakkara, T., and Shadmehr, R. (2002). A real-time state predictor in motor control: study of saccadic eye movements during unseen reaching movements. J. Neurosci. 22, 7721–7729.

Benson, B.L., Anguera, J.A., and Seidler, R.D. (2011). A spatial explicit strategy reduces error but interferes with sensorimotor adaptation. J. Neurophysiol. 105, 2843–2851.

de Brouwer, A.J., Albaghdadi, M., Flanagan, J.R., and Gallivan, J.P. (2018). Using gaze behavior to parcellate the explicit and implicit contributions to visuomotor learning. J. Neurophysiol. 120, 1602–1615.

Forano, M., and Franklin, D.W. (2019). Timescales of motor memory formation in dual-adaptation. BioRxiv 698167.

Galea, J.M., Mallia, E., Rothwell, J., and Diedrichsen, J. (2015). The dissociable effects of punishment and reward on motor learning. Nat. Neurosci. 18, 597–602.

Ghilardi, M., Ghez, C., Dhawan, V., Moeller, J., Mentis, M., Nakamura, T., Antonini, A., and Eidelberg, D. (2000). Patterns of regional brain activation associated with different forms of motor learning. Brain Res. 871, 127–145.

Haar, S., Donchin, O., and Dinstein, I. (2015a). Dissociating Visual and Motor Directional Selectivity Using Visuomotor Adaptation. J. Neurosci. 35, 6813–6821.

Haar, S., Givon-Mayo, R., Barmack, N.H., Yakhnitsa, V., and Donchin, O. (2015b). Spontaneous Activity Does Not Predict Morphological Type in Cerebellar Interneurons. J. Neurosci. 35, 1432–1442.

Haith, A.M., and Krakauer, J.W. (2018). The multiple effects of practice: skill, habit and reduced cognitive load. Curr. Opin. Behav. Sci. 20, 196–201.

Hegele, M., and Heuer, H. (2010). Implicit and explicit components of dual adaptation to visuomotor rotations. Conscious. Cogn. 19, 906–917.

Kim, H.E., Morehead, J.R., Parvin, D.E., Moazzezi, R., and Ivry, R.B. (2018). Invariant errors reveal limitations in motor correction rather than constraints on error sensitivity. Commun. Biol. 1, 19.

Kim, H.E., Parvin, D.E., and Ivry, R.B. (2019). The influence of task outcome on implicit motor learning. Elife 8.

Krakauer, J.W., Ghilardi, M., and Ghez, C. (1999). Independent learning of internal models for kinematic and dynamic control of reaching. Nat. Neurosci. 2, 1026–1031.

Leow, L.-A., Gunn, R., Marinovic, W., and Carroll, T.J. (2017). Estimating the implicit component of visuomotor rotation learning by constraining movement preparation time. J. Neurophysiol. 118, 666–676.

Mazzoni, P., and Krakauer, J. (2006). An Implicit Plan Overrides an Explicit Strategy during Visuomotor Adaptation. J. Neurosci. 26, 3642–3645.

McDougle, S.D., and Taylor, J.A. (2019). Dissociable cognitive strategies for sensorimotor learning. Nat. Commun. 10, 40.

McDougle, S.D., Bond, K.M., and Taylor, J.A. (2015). Explicit and Implicit Processes Constitute the Fast and Slow Processes of Sensorimotor Learning. J. Neurosci. 35, 9568–9579.

Morehead, J.R., Taylor, J.A., Parvin, D.E., and Ivry, R.B. (2017). Characteristics of Implicit Sensorimotor Adaptation Revealed by Task-irrelevant Clamped Feedback. J. Cogn. Neurosci. 29, 1061–1074.

Rabe, K., Livne, O., Gizewski, E.R., Aurich, V., Beck, A., Timmann, D., and Donchin, O. (2009). Adaptation to visuomotor rotation and force field perturbation is correlated to different brain areas in patients with cerebellar degeneration. J. Neurophysiol. 101, 1961–1971.

Rand, M.K., and Rentsch, S. (2016). Eye-Hand Coordination during Visuomotor Adaptation with Different Rotation Angles: Effects of Terminal Visual Feedback. PLoS One 11, e0164602.

Rentsch, S., and Rand, M.K. (2014). Eye-hand coordination during visuomotor adaptation with different rotation angles. PLoS One 9.

Smith, M.A., Ghazizadeh, A., and Shadmehr, R. (2006). Interacting adaptive processes with different timescales underlie short-term motor learning. PLoS Biol. 4, e179.

Taylor, J.A., and Ivry, R.B. (2011). Flexible cognitive strategies during motor learning. PLoS Comput. Biol. 7.

Taylor, J.A., and Ivry, R.B. (2013). Context-dependent generalization. Front. Hum. Neurosci. 7, 1–14.

Taylor, J. a, Krakauer, J.W., and Ivry, R.B. (2014). Explicit and implicit contributions to learning in a sensorimotor adaptation task. J. Neurosci. 34, 3023–3032.

Tipping, M.E., and Bishop, C.M. (1999). Probabilistic Principal Component Analysis. J. R. Stat. Soc. Ser. B (Statistical Methodol. 61, 611–622.

Werner, S., Van Aken, B.C., Hulst, T., Frens, M.A., Van Der Geest, J.N., Strüder, H.K., and Donchin, O. (2015). Awareness of sensorimotor adaptation to visual rotations of different size. PLoS One 10, 1–18.

Wong, A.L., Marvel, C.L., Taylor, J.A., and Krakauer, J.W. (2019). Can patients with cerebellar disease switch learning mechanisms to reduce their adaptation deficits? Brain 142, 662–673.

Zhang, Y., Wang, W., Zhang, X., and Li, Y. (2008). A cluster validity index for fuzzy clustering. Inf. Sci. (Ny). 178, 1205–1218.

